# The crucial role of intercellular calcium wave propagation triggered by influenza A virus in promoting infection

**DOI:** 10.1101/2024.12.20.629723

**Authors:** Fumiya Kozawa, Tomokazu Tamura, Naoki Takahashi, Taishi Kakizuka, Taro Ichimura, Rumi Shimada, Yasuyuki Hashimoto, Hironoshin Onituska, Sayaka Kashiwagi, Tomoko Kamasaki, Maho Amano, Takeharu Nagai, Takasuke Fukuhara, Yoichiro Fujioka, Yusuke Ohba

## Abstract

Influenza A viruses (IAVs) initially infect a few host cells, and the infection subsequently spreads to neighboring cells. However, the molecular mechanisms underlying this expansion remain unclear. Here, we show that IAV infection upregulates the frequency of intercellular calcium wave propagations (iCWPs) that mediate the spread of IAVs. ADP released from initially infected cells mediated iCWPs via the P2Y_1_ receptor. The generation of iCWPs and spread of viral infection were inhibited by a P2Y_1_ antagonist. Enhanced endocytosis in the surrounding cells that received ADP signaling upregulated viral entry. Expression of IAV matrix protein 2 (M2) in initially infected cells triggered iCWPs through ADP diffusion, thereby increasing infection, whereas an ion permeability-deficient mutation of M2 or inhibition of its ion channel activity suppressed iCWPs. Therefore, intercellular calcium signaling is essential for early expansion and potential establishment of IAV infection, which offers a promising target for IAV prophylaxis.

## Introduction

Influenza A virus (IAV), a member of the Orthomyxoviridae family, primarily infects mammals and waterfowl and causes acute respiratory tract infections. Despite the development of effective antiviral drugs and vaccines, it evokes annual seasonal epidemics and occasionally global pandemics with the emergence of mutant or reassortant strains (Uyeki *et al*, 2022; Steel & Lowen, 2014; Krammer *et al*, 2018; Herfst *et al*, 2012). IAVs have an eight-segment, single-stranded, and negative-sense RNA genome helically wrapped around nucleocapsid protein (NP). An RNA-dependent RNA polymerase (P) complex consisting of PA, PB1, and PB2 is bound to each end of the segment (Herfst *et al*, 2012). These viral ribonucleoproteins are enveloped by a matrix protein 1 (M1)-lined lipid bilayer to form the core. The outer surface of IAV particles consists of two spike proteins, hemagglutinin (HA) and neuraminidase (NA), and a relatively small number of matrix protein 2 (M2) (Neumann *et al*, 2009; Fontana & Steven, 2015).

Influenza A virus adsorption to the host cell surface is mediated by the binding of HA to sialylated cell surface proteins, including epidermal growth factor receptor, mannose receptor, and voltage-dependent calcium channel Ca_v_1.2 (Upham *et al*, 2010; Eierhoff *et al*, 2010; Fujioka *et al*, 2018; Kumlin *et al*, 2008). The binding of HA to Ca_v_1.2 results in an increase in intracellular Ca^2+^ concentration and the activation of various signaling pathways, which in turn promotes both clathrin-mediated endocytosis and macropinocytosis (Fujioka *et al*, 2013, 2018, 2011; de Vries *et al*, 2011), facilitating the entry of viral particles. Ca^2+^ dynamics is essential in infections by viruses other than IAVs. For example, hepatitis B virus (HBV) X protein (HBx) activates cytosolic Ca^2+^-dependent proline-rich tyrosine kinase-2 and promotes Ca^2+^ release from mitochondria, increasing intracellular Ca^2+^ concentration and subsequent replication of HBV DNA (Bouchard *et al*, 2001). The replication of respiratory syncytial virus is promoted via the induction of endoplasmic reticulum stress by nonstructural protein 2 and subsequent store-operated calcium entry (Cervantes-Ortiz *et al*, 2016). Furthermore, Ca^2+^ dynamics are involved in the entry of porcine reproductive and respiratory syndrome virus, rubella virus, severe fever with thrombocytopenia syndrome virus, and Ebola virus into their host cells, although the detailed molecular mechanisms remain to be determined (Diao *et al*, 2023; Li *et al*, 2019; Dubé *et al*, 2016; Nathan *et al*, 2020).

Neuraminidase is an exosialidase, which cleaves the α-ketoside bond between sialic acid and an adjacent sugar residue. During infection, it removes sialic acids from the glycoproteins of both host cells and viruses, thereby inhibiting reassociation of progeny virus particles with and aggregation on the plasma membrane via HA–sialic acid interactions. Finally, progeny viruses can bud off from infected cells and attach to noninfected cells, facilitating the spread of infection (McAuley *et al*, 2019; Gamblin & Skehel, 2010).

The third IAV particle surface protein, M2, is a virus-encoded ion channel, viroporin. Viroporin is a small transmembrane protein encoded by various viruses (Xia *et al*, 2022; Tomar *et al*, 2019). It contains one or two transmembrane domains and oligomerizes to form a tetrameric aqueous pore, enabling viral particles and host cell membranes to become permeable to ions, including Ca^2+^, H^+^, and K^+^. M2 protein of IAV facilitates viral infection through the following two mechanisms: First, in virus particles, it promotes fusion between the virus and the endosomal membrane by acidifying and maturing the endosomes during the entry process. Second, in virus-infected cells, M2 localizes to mitochondria and increases Ca^2+^ concentrations in both the cytosol and mitochondria, thereby positively regulating mitochondrial antiviral signaling-mediated innate immunity (Wang *et al*, 2019; Peng *et al*, 2021).

In this study, we discovered a third function of M2. The expression of M2 in infected cells resulted in a transient increase in intracellular Ca^2+^ concentration and ADP secretion, which propagated the wave of Ca^2+^ elevation from infected cells to the surrounding noninfected cells via the ADP receptor P2Y_1_. The transmission of calcium waves via cell-to- cell communication promoted viral internalization through activating endocytosis. This led to increased susceptibility to infection of noninfected cells surrounding infected cells.

## Results

### Transient increases in intracellular Ca^2+^ concentration propagate within cells surrounding infected cell

To analyze cell-to-cell Ca^2+^ dynamics within living cells, we established Madin–Darby canine kidney (MDCK) cells stably expressing the orange fluorescent variant of a genetically encoded Ca^2+^ indicator for optical imaging, O-GECO (MDCK/O-GECO), and cultured the cells in Matrigel to form monolayers. Under a conventional epifluorescence microscope, cells were exposed to an IAV strain A/Puerto Rico/8/34 (H1N1; PR8) at a multiplicity of infection (MOI) of 0.01 plaque-forming unit per cell (PFU/cell). Twelve-h post-infection (hpi), the increased intracellular Ca^2+^ concentration that originated from the initial firing cell was evident as it propagated outwards in concentric circle-like ripples to the surrounding cells (**Figs. 1A**,**B**). This phenomenon was termed intercellular calcium wave propagation (iCWP) because of its similarity to the calcium dynamics in thymic epithelial cells (Nihei *et al*, 2003). The frequency of iCWPs increased after 12 hpi, reached a maximum at 24 hpi, and returned to the baseline levels after 36 hpi (**Fig. 1C**). Given that virus-infected cell population rapidly increased between 12 and 36 hpi (**Fig. EV1A**), we hypothesized that a link exists between iCWPs and the spread of infection. Additionally, a “post” immunofluorescence assay using antibodies against NP, performed after observing the calcium dynamics, showed the existence of infected cells at the epicenter of propagation (**Fig. EV1B**). These iCWPs were also observed in the monolayer of human airway epithelial-derived BEAS-2B cells (**Fig. 1D**).

**Figure 1.**
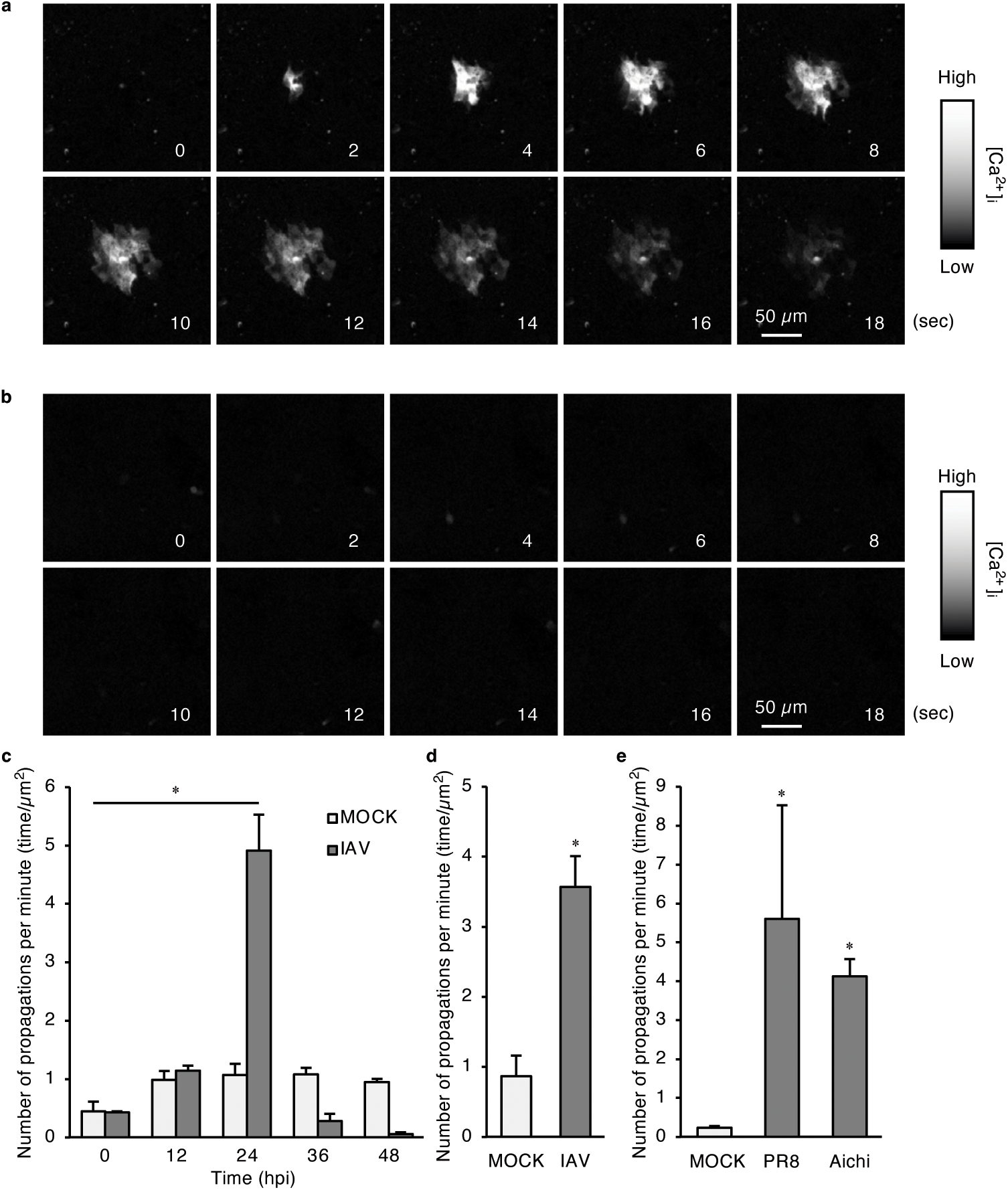
Influenza A virus (IAV) infection triggers intercellular calcium wave propagations (iCWPs) (**a**, **b**) MDCK cells stably expressing O-GECO (MDCK/O-GECO cells) were cultured in Matrigel for 5 days. After forming an epithelial monolayer, cells were exposed to influenza A virus A/Puerto Rico/8/34 (H1N1; PR8) at a multiplicity of infection (MOI) of 0.01 plaque- forming unit per cell (PFU/cell) (**a**) or phosphate-buffered saline (PBS) (**b**). After 24 h post- infection (hpi), cells were subjected to fluorescence microscopic observation. Representative time-lapse images at the time points indicated at the bottom right are shown. Scale bar, 50 μm. (**c**) MDCK/O-GECO cells were exposed to PR8 (IAV) or MOCK-infected as in (**a**) and observed under a fluorescence microscope. The number of iCWPs in 5 min was counted every 12 h. Data are represented as mean + standard error of the mean (SEM) (*n* = 3). \**p* < 0.0001 vs. MOCK-infected cells [one-way analysis of variance (ANOVA) with Dunnett’s post hoc test]. (**d**) BEAS-2B cells stably expressing O-GECO were cultured in Matrigel for 7 days and infected with PR8 at an MOI of 0.1 PFU/cell. At 24 hpi, cells were observed using a fluorescence microscope, and the number of iCWPs was counted. Data are mean + SEM (n = 3). \**p* = 0.0046 vs. MOCK-infected cells (Student’s *t*-test). (**e**) MDCK/O-GECO cells were infected with PR8, or A/Aichi/2/68 (H3N2; Aichi) at an MOI of 0.01 PFU/cell or mock-infected (MOCK). At 23 hpi, cells were observed using a fluorescence microscope to determine the frequency of iCWPs. \**p* < 0.0001 vs. MOCK- infected cells (one-way ANOVA with Dunnett’s post hoc test).

Moreover, IAV strains other than PR8 and A/Aichi/2/68 (H3N2; Aichi) also triggered iCWPs. Therefore, iCWPs induced by infection were not strain-specific (**Fig. 1E**).

Next, cells were monitored using a trans-scale imaging system (AMATERAS-1, a multiscale/modal analytical tool for every rare activity in singularity) (Ichimura *et al*, 2021). This imaging system allows simultaneous observation of approximately 1.0 × 10^6^ cells and enables large-scale quantitative analysis of rare biological events. We noticed the occurrence of iCWPs in uninfected monolayers with a relatively large propagation area, albeit at a relatively low frequency (**Figs. 2A**-**D**). This frequency increased 3-fold after IAV infection at 24 hpi (**Fig. 2E**). Furthermore, mapping the frequency of iCWP in a field of view (200 μm^2^) over 10 min identified the sites where wave propagation occurred multiple times in IAV-infected cells (**Figs. 2F**,**G**). Consequently, large-scale quantitative analysis suggested the presence of hotspots for the origin of IAV-induced iCWPs.

**Figure 2.**
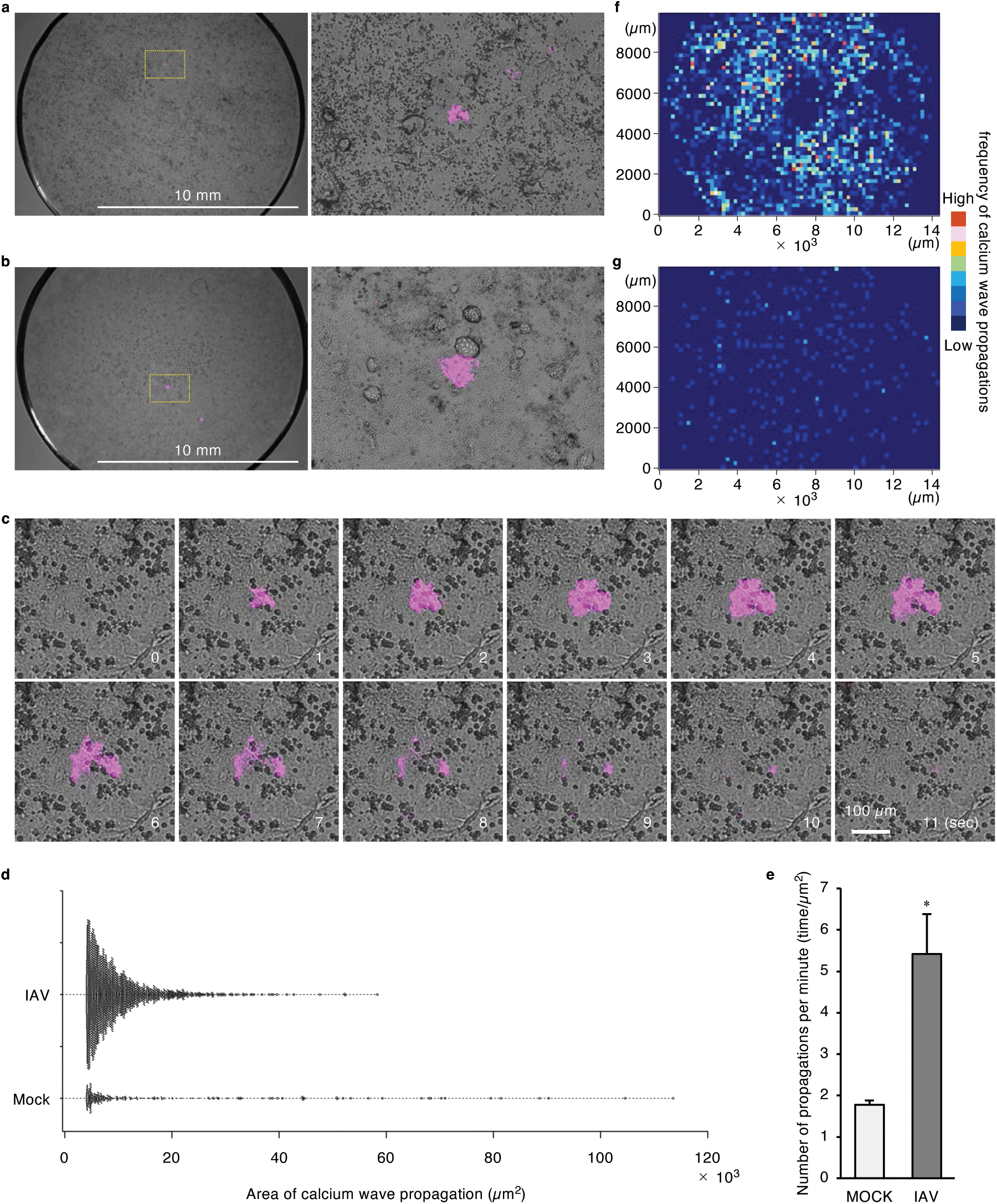
iCWPs triggered by virus infection are repeatedly generated in specific regions. (**a**–**e**) Epithelial monolayers of MDCK/O-GECO cells were exposed to PR8 at an MOI of 0.01 PFU/cell (**a**) or PBS (**b**). At 24 hpi, cells were observed using an AMATERAS-1 system. Representative low-magnification merged images of transmission (gray) and O-GECO (magenta) (left), and high-magnification images of yellow squares therein (right) are shown (**a**). Scale bar, 10 mm. Representative time-lapse images (**a**, right) at the time points indicated in the bottom right are shown (**c**). Scale bar, 100 μm. The area (**d**) and frequency (**e**) of iCWPs were quantified and plotted. Data are represented as mean + SEM (n = 3). \**p* = 0.0064 vs. MOCK-infected cells (Student’s *t*-test). (**f,g**) The field of view was segmented into 200 × 200 µm square compartments, and the number of iCWPs observed in 10 min in each area was mapped using color hues according to the lookup table for infected (**f**) and control cells (**g**).

### Influenza M2 protein is responsible for initiating iCWPs

To determine the viral protein(s) responsible for initiating iCWPs, we observed the calcium dynamics in a monolayer of MDCK/O-GECO cells that had been transfected with the expression vectors for HA and M2 of the PR8 strain, both of which have been implicated in regulating intracellular Ca^2+^ concentration (Wang *et al*, 2019; Fujioka *et al*, 2018). The expression of these proteins was monitored by bicistronic expression of enhanced green fluorescent protein (EGFP) mediated by an internal ribosome entry site (IRES). iCWPs appeared to spread from cells expressing M2 (**Fig. 3A**), and their frequency was significantly higher than that in control vector-transfected or HA-expressing cells (**Fig. 3B**). Given that treatment with the M2 proton channel inhibitor, amantadine (Pinto *et al*, 1997), reduced the number of iCWPs induced by M2 in the amantadine-sensitive Aichi strain (**Fig. 3C**), channel pore function was required for the generation of iCWPs. Notably, the introduction of an ion permeability-deficient mutant of M2 (A27F) (Balannik *et al*, 2010) failed to induce iCWPs (**Fig. 3D**), although it localized to mitochondria labeled with TOM5, similar to the wild-type form (**Fig. EV2**). These results indicated that ion channel activity of M2 in mitochondria might be required for inducing wave propagation. Therefore, we investigated the effect of inhibitors of the protein regulating mitochondrial ion dynamics on the induction of M2- and IAV-induced iCWPs. Treatment with venetoclax, a B-cell lymphoma 2 (Bcl-2) family protein inhibitor, significantly suppressed iCWPs induced by M2 expression, whereas treatment with an Na^+^/Ca^2+^ exchanger inhibitor (CGP37157 [CGP]) and a mitochondrial permeability transition pore inhibitor (cyclosporin A [CsA]) failed to do so (**Fig. 3E**). Furthermore, venetoclax suppressed IAV-induced iCWPs (**Fig. 3F**). Therefore, Bcl-2 family proteins are crucial for generating M2-mediated iCWPs following IAV infection.

**Figure 3.**
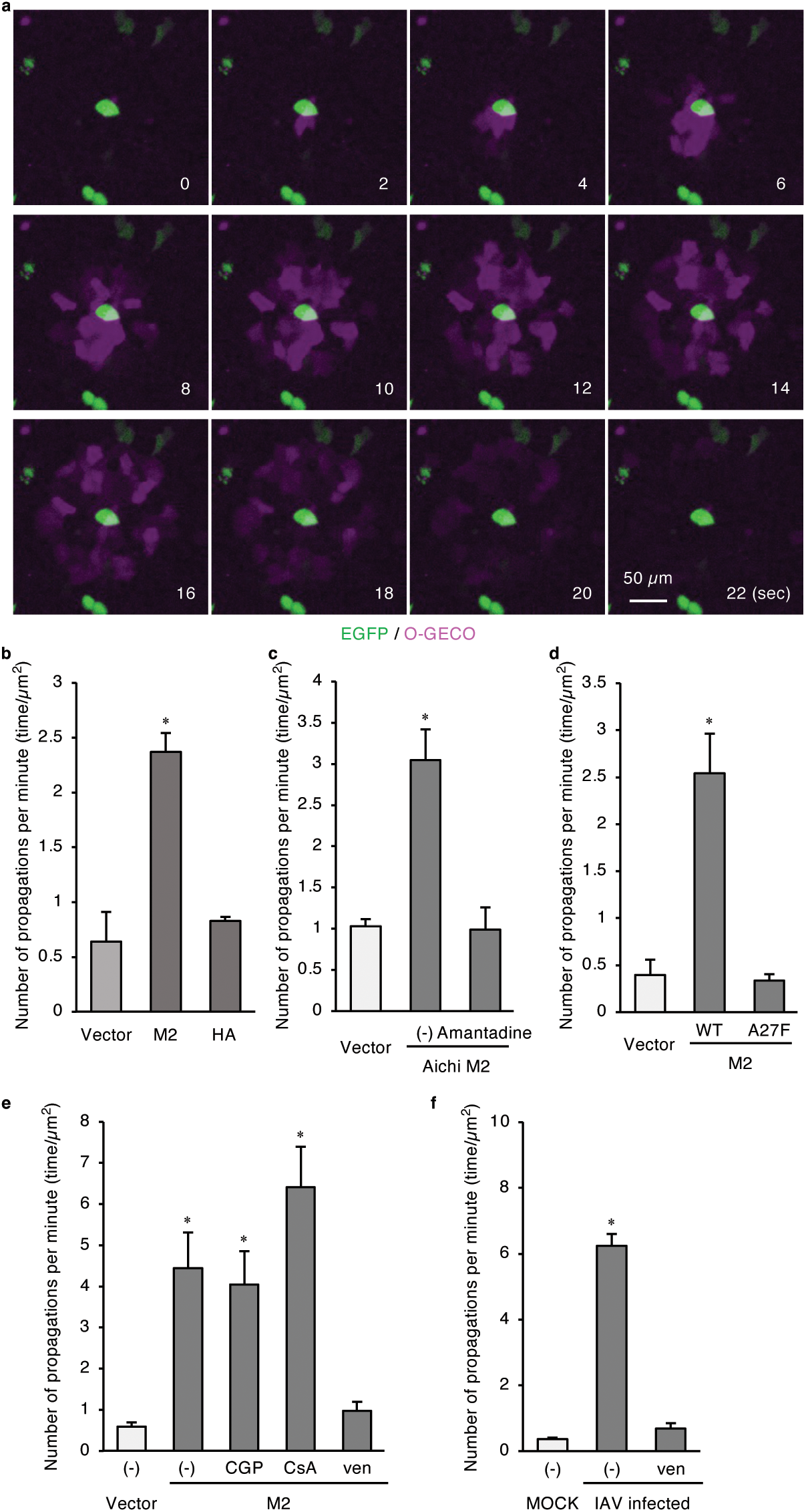
IAV protein M2 provokes iCWPs via its ion channel activity. (**a, b**) MDCK/O-GECO cells were transfected with the control vector, an expression vector for PR8 M2, or that for PR8 HA. After 24 h, cells were observed using a fluorescence microscope, and the number of iCWPs was quantified. Representative time-lapse, merged images of M2 (green) and O-GECO (magenta) of the cell population around PR8 M2- expressing cells, monitored by bicistronic expression of enhanced green fluorescent protein (EGFP), at the time points indicated at the bottom right are shown (**a**). Scale bar, 50 μm. The frequency of iCWPs was quantified and plotted (**b**). Data are presented as mean + SEM of three independent experiments. *p* = 0.0011 (one-way ANOVA with Dunnett’s post hoc test). (**c**) MDCK/O-GECO cells transfected with a control vector or an expression vector for Aichi M2 were treated with amantadine for 1 h or left untreated ([-]). The frequency of iCWPs was quantified using an epifluorescence microscope and plotted as in (**b**). Data are presented as mean + SEM of three independent experiments. *p* = 0.0001 (one-way ANOVA with Dunnett’s post hoc test). (**d**) MDCK/O-GECO cells expressing a wild-type (WT) or a mutant (A27F) form of PR8 M2 were observed using a fluorescence microscope to determine the frequency of iCWPs. A respective empty vector (Vector) was used as a negative control. Data are presented as mean + SEM of three independent experiments. *p* = 0.0022 (one-way ANOVA with Dunnett’s post hoc test). (**e, f**) MDCK/O-GECO cells expressing M2 (**e**) or cells infected with PR8 at an MOI of 0.01 PFU/cell (**f**, IAV infected) were treated with the indicated inhibitors for 1 h or left untreated [(-)]. Respective empty vectors (**e**, Vector) or mock-infected cells (**f**, MOCK) were used as negative controls. The number of iCWPs was quantified and plotted as in (**b**). Data are represented as mean + SEM of three independent experiments. *p* < 0.01 (one-way ANOVA with Dunnett’s post hoc test).

### ADP–P2Y_1_ signaling mediates iCWPs

Given that the propagation rates of calcium waves in IAV-infected cells were comparable to those in M2-expressing cells (**Fig. EV3**), we hypothesized that these propagations might be transmitted from cell to cell through a common mechanism. Therefore, we aimed to elucidate the mediators of propagation. Because ATP/ADP signaling and gap junctions mediate iCWPs (Nihei *et al*, 2003; Justet *et al*, 2016; Jiang *et al*, 2017), we investigated the effect of ATP/ADP diphosphohydrolases (Apyrase VI and VII) and a gap-junction inhibitor 18β- glycyrrhetinic acid 2K on the generation of propagations. Apyrase VII, which catalyzes the hydrolysis of ADP to AMP (Handa & Guidotti, 1996; Matchkov *et al*, 2004), significantly suppressed IAV-induced iCWPs, whereas Apyrase Ⅵ, which hydrolyzes ATP, and 18β-glycyrrhetinic acid 2K did not (**Fig. 4A**). Therefore, ADP secreted from infected cells may be critical in mediating iCWPs. Moreover, the addition of caged-ADP to the culture and subsequent irradiation to ultraviolet (UV) light increased the fluorescence intensity of O-GECO in MDCK/O-GECO cells, indicating that the intracellular Ca^2+^ concentration was increased by extracellular ADP (**Fig. EV4A**).

**Figure 4.**
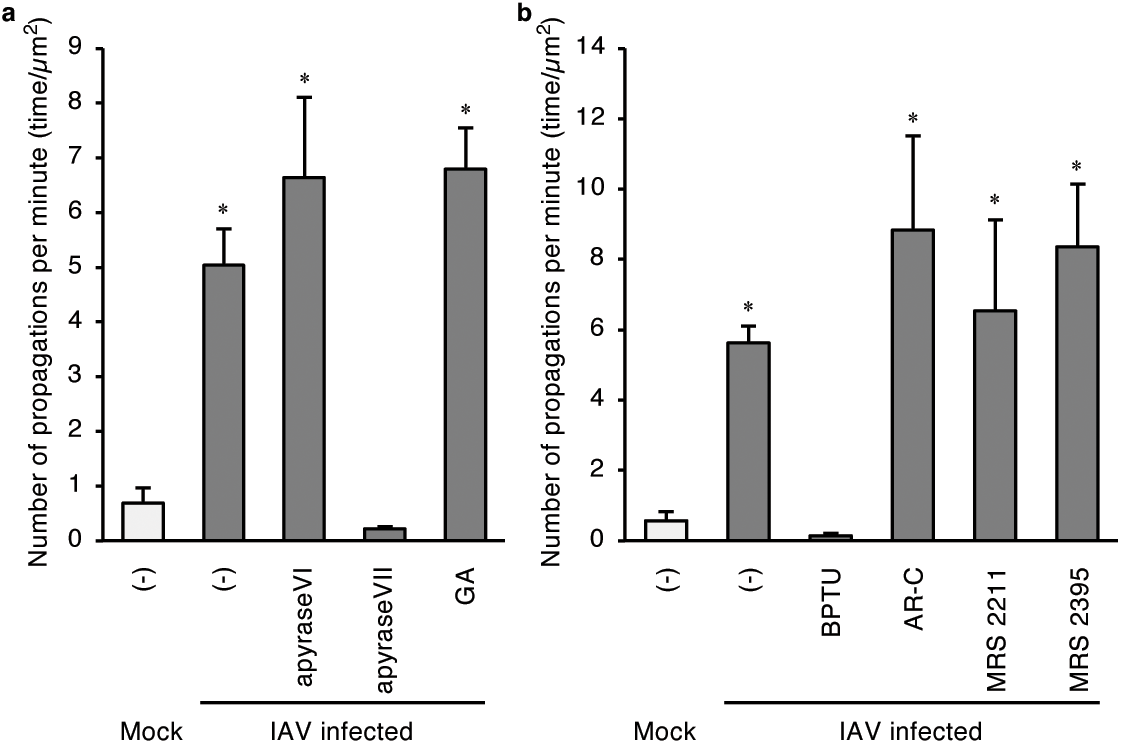
ADP–P2Y_1_ paracrine signaling mediates iCWPs. (**a, b**) MDCK/O-GECO cells were infected with PR8 at an MOI of 0.01 PFU/cell (IAV infected) or mock-infected. After 23 hpi, cells were treated with inhibitors indicated at the bottom [apyrase VI, apyrase VII, or 18β- glycyrrhetinic acid 2K (GA) (**a**); 1-(2-(2- *tert*- butylphenoxy)pyridin-3-yl)-3-(4-(trifluoromethoxy)phenyl)urea (BPTU), ARC-118925XX (AR-C), MRS2211, or MRS2395 (**b**)] for 1 h, or left untreated [(-)]. Cells were observed using a fluorescence microscope to determine the frequency of iCWPs. Data are represented as mean + SEM of three independent experiments. *p* < 0.05 (one-way ANOVA with Dunnett’s post hoc test).

The expression of the wild-type form of M2 resulted in a sustained increase in intracellular Ca^2+^ concentration in MDCK cells, whereas expression of the ion channel activity-deficient mutant form of M2, which could not induce iCWPs (**Fig. 3D**), failed to do so (**Fig. EV4B**). These results suggested that ADP was released from infected cells in a manner dependent on M2 expression and its pore function, as a sustained increase in intracellular Ca^2+^ concentration promotes ADP release from cells (Elaïb *et al*, 2016). Treatment with venetoclax abrogated the M2-dependent but not the ADP-dependent, increase in intracellular Ca^2+^ concentration (**Figs. EV4A**,**B**), indicating that Bcl-2 family proteins play a role in M2- and IAV-driven iCWPs in infected cells (ADP release) rather than in the surrounding cells.

To determine the cognate ADP receptor in iCWPs, we treated cells with inhibitors of the ADP receptors P2Y_1_, P2Y_2_, P2Y_12_, and P2Y_13_ (Wirsching *et al*, 2020; Maynard & Sfanos, 2022) for 1 h at 24 hpi. The number of iCWPs decreased following the treatment with the P2Y_1_ inhibitor 1-(2-(2- *tert*-butylphenoxy)pyridin-3-yl)-3-(4-(trifluoromethoxy)phenyl)urea (BPTU) at a noncytotoxic concentration (10 µM, **Fig. EV4C**). In contrast, inhibitors of P2Y_2_, P2Y_12_, and P2Y_13_ (AR-C 118925XX, MRS 2395, and MRS 2211, respectively) failed to do so (**Fig. 4B**). BPTU treatment also decreased the number of M2-induced iCWPs (**Fig. EV4D**). To exclude the possibility that progeny virus particles mediate iCWPs, we evaluated cell-to-cell Ca^2+^ dynamics in IAV-infected cells in the presence of sialidase and voltage-dependent calcium channel inhibitor diltiazem, which block virus adsorption and entry into host cells, respectively (Fujioka *et al*, 2018). These treatments did not alter the number of iCWPs (**Fig. EV4E**), suggesting that ADP, but not the progeny viruses, mediate wave propagation.

### iCWPs accelerate virus infection

To assess whether IAV-induced iCWPs promote the production of progeny viruses, we evaluated the effects of propagation inhibitors (**Figs. 4A**,**B**) on viral replication. Treatment with Apyrase VII and BPTU significantly suppressed the production of progeny viruses in the culture medium, as quantified by immunofluorescence using anti-NP antibodies (**Fig. 5A**) and plaque assay (**Fig. 5B**). In contrast, such inhibitory effects were not observed upon the treatment with Apyrase VI, AR-C 118925XX, and 18β-glycyrrhetinic acid 2K (**Figs. 5A**,**B**; see also **Figs. 3A**,**B**). The distribution pattern of NP-positive cells in the control was characterized by island-like cell cluster formation, whereas infected cells in the BPTU-treated samples were solitary and scattered (**Fig. EV5A**). Quantitative analysis revealed that the distribution of intercellular distances between IAV-infected cells shifted towards relatively small distances only for the BPTU-untreated samples, compared to the calculated values for randomly arranged cells (**Fig. EV5B**). These findings support the hypothesis that infection is promoted in areas where calcium waves propagate. We have previously reported that elevated intracellular Ca^2+^ concentrations activate endocytosis and promote viral incorporation (Fujioka *et al*, 2013). Therefore, we examined whether endocytosis was activated by iCWPs and found that dextran uptake by cells adjacent to the M2-expressing cells increased compared to cells surrounding the control GFP-expressing cells (**Fig. EV5C**). This suggests that ADP–P2Y_1_ signaling-mediated iCWPs from infected cells promote endocytosis and subsequent IAV infection in surrounding noninfected cells.

**Figure 5.**
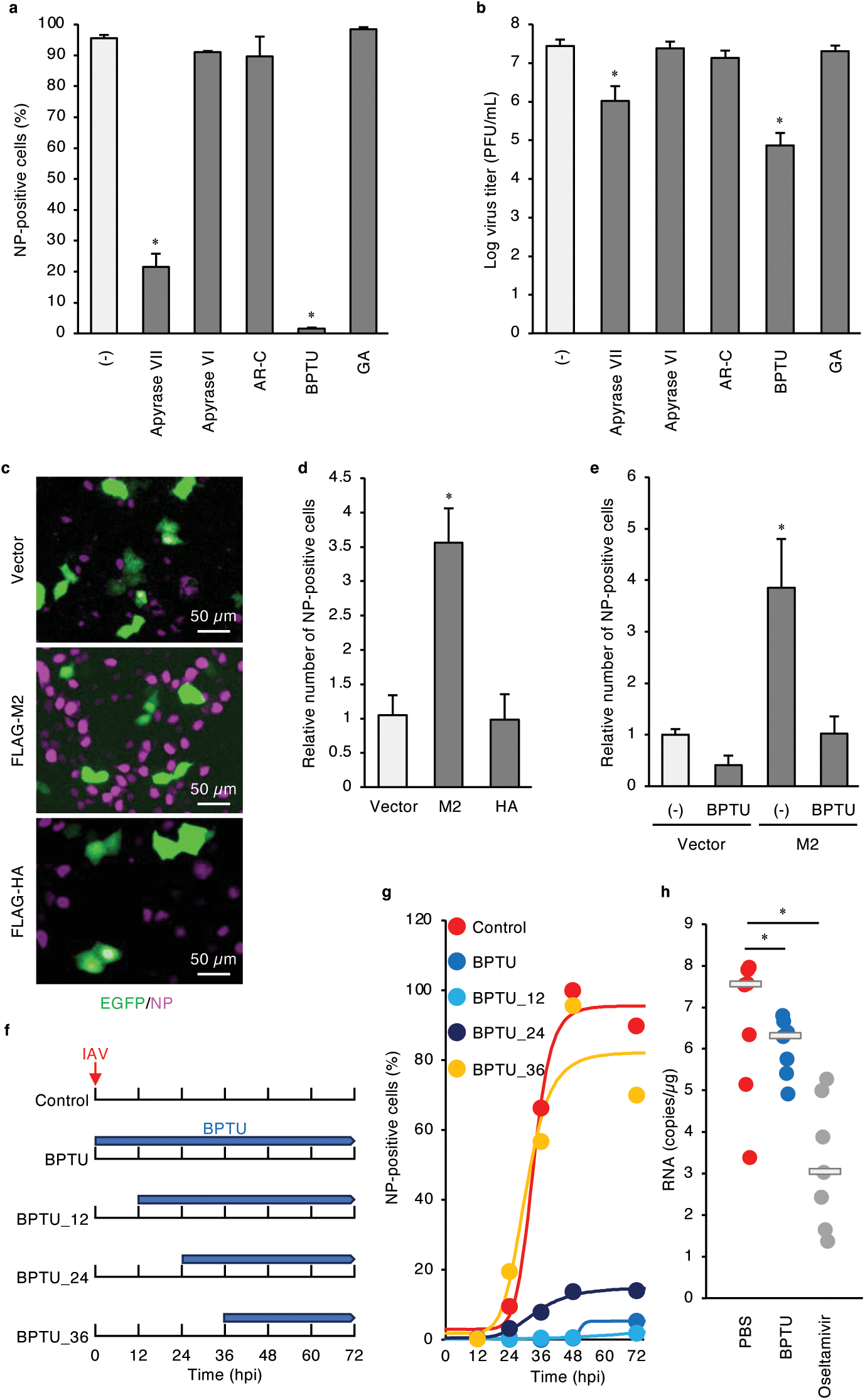
The P2Y_1_ antagonist BPTU inhibits IAV infection. (**a, b**) Epithelial monolayers of MDCK cells were pretreated with the indicated agents for 1 h or left untreated ([-]), and exposed to PR8 at an MOI of 0.01 PFU/cell. To evaluate virus replication, the culture medium was collected at 48 hpi, and the number of progeny viruses was determined by a plaque assay (**a**) and an immunofluorescence-based infection assay using antibodies against nucleocapsid protein (NP) (**b**). The virus titer (**a**) and percentage of NP- positive cells (**b**) were plotted. Data are presented as mean + SEM of three independent experiments. *p* < 0.05 (one-way ANOVA with Dunnett’s post hoc test). (**c, d**) MDCK cells were transfected with the control vector, pCXN2-FLAG-PR8 M2-IRES- EGFP, or pCXN2-FLAG-PR8 HA-IRES-EGFP. After 24 h, cells were exposed to PR8 at an MOI of 0.2 PFU/cell for 4 h, fixed, and subjected to an immunofluorescence-based infection assay. Representative merged images of bicistronically expressed EGFP (green) and NP (magenta) are shown (**c**). Scale bar, 50 µm. The percentage of NP-positive cells was plotted (**d**). Data are presented as mean + SEM of three independent experiments. *p* = 0.0007 (one- way ANOVA with Dunnett’s post hoc test). (**e**) MDCK cells transfected with a control vector or an expression vector for FLAG-PR8 M2 were treated with dimethyl sulfoxide (DMSO) ([-]) or BPTU and infected with PR8 at an MOI of 0.2 PFU/cell. The number of infected cells at 4 hpi was determined by an immunofluorescence-based infection assay. Data are presented as mean + SEM of three independent experiments. *p* = 0.011 (one-way ANOVA with Turkey’s HSD post hoc test). (**f**) Treatment regimen in a time-of-drug-addition assay. (**g**) MDCK cells cultured in Matrigel were exposed to PR8 at an MOI of 0.01 PFU/cell and treated with BPTU according to the regimen shown in (**F**). The cultured medium for each set was collected at 12, 24, 36, 48, and 72 hpi. The number of progeny viruses in the culture medium was determined by an immunofluorescence-based infection assay and plotted. The solid lines are the Hill curves generated by fitting each data set. (**h**) BALB/c mice (4-week-old, male, *n* = 7) per group were intraperitoneally administrated BPTU, oseltamivir, or PBS and then intranasally inoculated with PR8. The lungs were collected at 2 days post-infection, and viral RNA copy number was determined by quantitative PCR (qPCR). Data are presented as mean + SEM from an experiment. *p* < 0.03 (one-way ANOVA with Turkey’s HSD post hoc test).

Next, to directly confirm the involvement of M2 in promoting IAV infection by iCWP, MDCK epithelial monolayers containing a small number of M2-expressing cells were formed and exposed to viruses. NP-positive virus-infected cells accumulated around M2- expressing cells (**Fig. 5C**), which were the epicenters of iCWP, and their numbers were significantly higher than those in other areas or in the controls (**Fig. 5D**). In contrast, HA expression did not exhibit this phenomenon (**Figs 5C**,**D**). The infection-promoting effect of M2 was almost entirely cancelled by BPTU treatment (**Fig. 5E**).

In addition, to implicate iCWP in the expansion of viral infection, BPTU was administered according to the regimen of a time-of-drug-addition assay (**Fig. 5F**) (Yan *et al*, 2022), and time-dependent changes in progeny virus production were quantified by an immunofluorescence-based virus infection assay using NP antibodies. When the treatment was started at an early stage of infection (i.e., until 12 hpi), the expansion of infection was almost completely inhibited (**Fig. 5G**). The expansion of infection was also attenuated when treatment was initiated after 24 hpi; however, this inhibitory effect was not observed after 36 hpi (**Fig. 5G**). Given that 24 hpi is the period when iCWPs most frequently emerge (**Fig. 1C**), and the number of infected cells exponentially increases (**Figs. 5G** and **1D**), this result strongly implicates iCWPs in the expansion of infected cells.

Finally, to investigate the impact of ADP–P2Y_1_ signaling in virus infection *in vivo*, we intraperitoneally pre-administered BPTU, antiviral oseltamivir, or phosphate-buffered saline to mice and evaluated whether the reagents could prevent viral spread in mice. Viral RNA loads in the lungs, as determined by quantitative polymerase chain reaction (qPCR), were lower in the BPTU- and oseltamivir-treated groups than in the control group (**Fig. 5H**). These results demonstrate that ADP–P2Y_1_ signaling is operative, and its inhibition can be a therapeutic target for IAV infection *in vivo*.

## Discussion

In this study, we demonstrate that IAVs induce iCWPs from an infected cell to the surrounding noninfected cells, thereby promoting endocytosis and subsequent viral infection. The expression of IAV M2 proteins in infected cells induces sustained elevation of intracellular Ca^2+^ concentration in a manner dependent on its ion channel function and Bcl-2 family proteins, eliciting ADP release. ADP–P2Y_1_ signaling mediates the propagation of calcium waves and expansion of IAV infection to the surrounding cells.

How does iCWP promote IAV infection? The first possibility is that iCWP increases intracellular Ca^2+^ concentration and upregulates endocytic activity of cells (Fujioka *et al*, 2018, 2013), thereby creating a “field” more susceptible to infection. Endocytosis was activated in cells surrounding M2-expressing cells. The second possibility is that elevated Ca^2+^ concentrations may promote replication of viral RNA; for example, hepatitis C and Zika viruses promote Ca^2+^ influx to translocate DEAD-box helicase 3 X-linked from the plasma membrane to the cytoplasm, which facilitates translation of viral RNA (Doñate-Macián *et al*, 2018). Another possibility may be the upregulation of egress of progeny particles. Budding of filoviruses (Ebola and Marburg) and arenaviruses (Lassa and Junín viruses) is controlled by intracellular Ca^2+^ concentrations (Nathan *et al*, 2020; Dewald *et al*, 2018; Han *et al*, 2015).

Regardless of other possibilities, iCWPs likely represent a sophisticated invasion strategy of viruses; a small number of vanguard virus particles take the initiative and establish a front- line base (the epicenter cell of propagation), from which a “field” susceptible to infection is generated by scouting neighboring cells using iCWPs.

Rotaviruses generate iCWPs in human intestinal enteroids (Chang-Graham *et al*, 2020). Unlike IAVs, the infection of which is upregulated by iCWPs, rotaviruses employ this phenomenon to elicit systemic inflammatory responses through increased mRNA expression of the inflammatory cytokine interleukin-1α in the enteroids. Given that the pathogenesis of severe influenza virus infection is owing to excessive immune response, determining whether IAV-induced iCWP has a role similar to that of rotavirus-induced infection is of prime interest.

iCWPs have been implicated in pathophysiological functions in various cellular contexts. For example, we have previously reported that calcium waves propagate from cancer cells to the surrounding normal cells, eliminating cancer cells from the epithelial monolayer (Takeuchi *et al*, 2020). During nerve injury, iCWP induced by ADP release from apoptotic neurons results in oriented migration of microglia (Jiang *et al*, 2017). Tissue wounding of the corneal endothelia triggers iCWPs to control cell proliferation during wound healing (Handly & Wollman, 2017). Such iCWPs in noninfected tissues are likely mediated by a mechanism exactly similar to that during infection.

Elevated intracellular calcium concentration in a cell is directly or indirectly transmitted to the neighboring cells. Direct transmission is mediated by gap junctions, through which Ca^2+^ is transported to the adjacent cells. In the latter case, secreted ATP or ADP diffuses and binds to the G_q_-coupled purinergic receptors on the neighboring cells, thereby increasing intracellular Ca^2+^ concentrations (Maynard & Sfanos, 2022; Wirsching *et al*, 2020). Among the ADP receptors, P2Y_1_ was identified as an essential factor in mediating iCWPs during IAV infection in this study because the P2Y_1_-specific antagonist BPTU suppressed propagation.

P2Y_1_ is ubiquitously expressed *in vivo*, including the respiratory tract, whereas the expression of P2Y_12_ and P2Y_13_ is restricted to the bone marrow and immune cells (Conroy *et al*, 2016). The fact that inhibitors of P2Y_12_ and P2Y_13_ receptors did not suppress iCWPs or IAV infection might be explained by low expression of these receptors in the cells used in this study. Notably, rotavirus also induces wave propagation through the expression of nonstructural protein 4, an endoplasmic reticulum-localized viroporin (Chang-Graham *et al*, 2020). Therefore, IAV- and rotavirus-induced iCWPs share a common underlying mechanism; both are triggered by viroporin expression and mediated by ADP–P2Y_1_ signaling (**Fig. EV4D**). Given that this mechanism also eliminates cancer cells, microglial migration, and wound healing of tissues (Takeuchi *et al*, 2020; Jiang *et al*, 2017; Handly & Wollman, 2017), viruses may cleverly hijack the host cell machinery to create an environment favorable for their infection.

The mechanism by which influenza viruses increase Ca^2+^ concentration in initially firing cells might be shared by other viruses; during HBV infection, the viral protein HBx locates Bcl-2 and Bcl-extra large (Bcl-xL) to mitochondria and induces sustained elevation of intracellular Ca^2+^ concentrations, thereby promoting replication of the viral genome and host cell death (Bouchard *et al*, 2001; Geng *et al*, 2012). Furthermore, the expression of Bcl-xL increases at 24 hpi of IAV infection (Lee *et al*, 2018). Notably, cytosolic Ca^2+^ concentration sustainably increased in M2-expressing (initial IAV-infected) cells, from which ADP was secreted and inhibited by the Bcl-2 family protein inhibitor venetoclax (**Fig. EV4B**). Given that venetoclax did not inhibit ADP-dependent elevation in Ca^2+^ concentrations (**Fig. EV4A**), the antiapoptotic Bcl-2 family proteins may contribute to ADP production/release in the epicentral cell of iCWP generation; however, they are not involved in propagation in surrounding cells.

Parasitic pathogenic viruses infect host cells by hijacking host cell factors and functions, regardless of the emergence of mutant or reassortant strains, which often provides potential targets for developing broad-spectrum therapies. Furthermore, iCWPs demonstrated in the present study were observed in multiple strains. The results presented here may help engineer a new class of potent therapeutic molecules, including P2Y_1_ inhibitors and antagonists, irrespective of viral serotypes.

## Methods

### Ethics statement

All experiments with mice were performed in accordance with the Science Council of Japan Guidelines for the Proper Conduct of Animal Experiments. The study protocol was approved by the Institutional Animal Care and Use Committee of National University Corporation, Hokkaido University (approval ID: 22-0105).

### Cell culture and transfection

Madin-Darby canine kidney (MDCK, CCL-34) and human bronchial epithelial (BEAS-2B, CRL-9609) cells were obtained from American Type Culture Collection (Manassas, VA, USA). Cells were maintained in a humidified atmosphere of 5% CO_2_ at 37 °C in Dulbecco’s modified Eagle’s medium (DMEM; Sigma-Aldrich, St. Louis, MO, USA) supplemented with 10% fetal bovine serum (Capricorn Scientific, Ebsdorfergrund, Germany). Expression plasmids (2.5 µg, unless otherwise specified) were introduced into MDCK, or BEAS-2B cells by transfection for 24 h using FuGENE HD (Promega, Madison, WI, USA) or Polyethylenimine “Max” (PEI-MAX; Polyscience, Warrington, PA, USA). The absence of *Mycoplasma* contamination was confirmed using a PCR Mycoplasma Test Kit (Takara Bio, Kusatsu, Japan). MDCK and BEAS-2B cells were cultured in Matrigel (BD Biosciences, Franklin Lakes, NJ, US) in a 35-mm-glass-base dish every 6 and 5 days, respectively, to prepare 3D monolayers.

### Purification of viruses

MDCK cells were infected with IAV strains A/Puerto Rico/8/34 (H1N1; PR8) or A/Aichi/2/68 (H3N2; Aichi) at a multiplicity of infection (MOI) of 0.0001 plaque-forming unit per cell (PFU/cell) for 48 h at 35 °C. The culture medium was centrifuged to remove cell debris, and the supernatant was collected. The viruses isolated by ultra-highspeed centrifugation were resuspended in phosphate-buffered saline (PBS) and stored at -80 °C until use. The titers of the infectious viruses in virus solution or culture media were determined using a plaque assay (Pleschka *et al*, 2001). In brief, a conditioned medium of virus-infected cells was inoculated into confluent MDCK cell monolayers in 12-well plates; after 1 h of inoculation, the supernatants were removed, and cells were incubated with minimal essential medium containing 1% Bacto-agar (Sigma-Aldrich) and trypsin (5 mg/ml, Sigma-Aldrich). Plaques were counted after incubation for 2 days at 35 °C under 5% CO_2_.

### Reagents and antibodies

Antibody against IAV nucleoprotein (NP, #GTX125989) was obtained from GeneTex (Irvine, CA, USA). The P2Y_2_ antagonist AR-C 118925XX (#216657-60-2) and P2Y_1_ antagonist 1-(2- (2- *tert*-butylphenoxy)pyridin-3-yl)-3-(4-(trifluoromethoxy)phenyl)urea (BPTU; BMS- 646786, #870544-59-5) were obtained from Tocris Bioscience (Bristol, UK). Apyrase grade VI (#A6410), apyrase grade VII (#A6535), and RITC-Dextran (#R9379) were purchased from Sigma-Aldrich, and 18β-glycyrrhetinic acid was obtained from Santa Cruz Biotechnology (Dallas, TX, USA). Hoechst 33342, carbocyanine dyes (DiO, DiI, and DiD), Alexa Fluor 488-labeled antibodies to mouse (#AB_2534069) or rabbit (#AB_2633280) immunoglobulin G, and Alexa Fluor 647-labeled antibodies to mouse (#AB_2535804) or rabbit (#AB_2535813) immunoglobulin G were obtained from Thermo Fisher Scientific (Carlsbad, CA, USA). Matrigel (Cat#256237) was obtained from BD Biosciences. CGP37157 (#HY-15754) and venetoclax (#GDC-0199) were obtained from MedChem Express (Monmouth, NJ, USA). Cyclosporin A (#C2408), oseltamivir phosphate (#GS-4104), NPE-caged-ADP, Horseradish peroxidase (HRP)-conjugated goat secondary antibodies against mouse/rabbit immunoglobulin G, and trypsin were obtained from Tokyo Chemical Industry (Tokyo, Japan), Cayman Chemical (Ann Arbor, MI, USA), Jena Bioscience (Malchin, Germany), Jackson ImmunoResearch Laboratories (West Grove, PA, USA), and FUJIFILM Wako (Osaka, Japan), respectively. MRS 2211 (2-[(2-chloro-5-nitrophenyl)azo]-5-hydroxy-6- methyl-3-[(phosphonooxy)methyl]-4-pyridinecarboxaldehyde disodium salt and MRS 2395 (2,2-dimethyl-propionic acid 3-(2-chloro-6-methylaminopurin-9- yl)-2-(2,2-dimethyl- propionyloxymethyl)-propyl ester) were obtained from Merck (Darmstadt, Germany).

### Plasmids

The expression vector for pCX4puro-CRY2-CRaf(Aoki *et al*, 2013) was generously provided by Dr. K. Aoki (National Institute for Basic Biology, Okazaki, Japan). The expression vector pCMV-O-GECO1 was purchased from Addgene (Cambridge, MA, USA). pCMV-O-GECO1 was digested with BamHI and NotI. The resulting DNA fragment (the coding sequence of O- GECO1) was subcloned between the BamHI and NotI sites of pCX4puro-CRY2-CRaf to obtain pCX4puro-O-GECO1. The coding sequences for HA and M2 were amplified from pCAGGS-HA and pCAGGS-M2 (Neumann *et al*, 1999), obtained from Dr. Y. Kawaoka, using the primer pairs HA_F and HA_R, and M2_F and M2_R, respectively. The resulting PCR products were digested with XhoI and NotI, and then subcloned between the XhoI and NotI sites of pCXN2-Flag-HRas-IRES-EGFP to obtain pCXN2-Flag-HA-IRES-EGFP and pCXN2-Flag-M2-IRES-EGFP. The expression vector for the A27F mutant of M2 was obtained by PCR-based mutagenesis using a QuickChange Site-directed mutagenesis kit (Agilent, Santa Clara, CA, USA) and with primers:

5’-GTGATCCTCTCTTTATTGCCGCAAATATC-3’ and

5’-GATATTTGCGGCAATAAAGAGAGGATCAC-3’.

### Generation of stable cell lines

pCX4puro-O-GECO1 was linearized with ScaI and introduced into BEAS-2B and MDCK cells by transfection using PEI MAX for 24 h. Cells were then cultured in DMEM supplemented with 10% fetal bovine serum and puromycin, and positive colonies were selected after 14 days. Cloned cells were selected using puromycin after every five passages.

### Fluorescence microscopy

Fluorescence imaging and data analysis were performed as previously described (Fujioka *et al*, 2019; Kashiwagi *et al*, 2019). Briefly, cells were placed in a stage-top incubation chamber maintained at 37 °C on a Nikon Eclipse T*i*2-E microscope (Nikon, Tokyo, Japan) equipped with a KINETIX22 scientific complementary metal oxide semiconductor (sCMOS) camera (Teledyne Photometrics, Tucson, AZ, USA), PlanApo 4×, 10×, 20×, or 60× objective lenses, a TI2-CTRE microscope controller (Nikon), and a TI2-S-SE-E motorized stage (Nikon). Cells were illuminated using an X-Cite Turbo system (Excelitas Technologies, Waltham, MA, USA). The sets of excitation and emission filters and dichroic mirrors adopted for this observation included GFP HQ (Nikon) for EGFP, Cy3 HQ (Nikon) for mCherry, or DAPI-U HQ (Nikon) for Hoechst 33342. For live-cell imaging, cells were incubated at 37 °C using a Chamlide incubator system (Live Cell Instrument, Seoul, Republic of Korea).

Confocal and super-resolution images were acquired using an X-Light V3 spinning disk confocal unit (CrestOptics, Roma, Italy) and DeepSIM (CrestOptics) for Eclipse Ti2 equipped with a Prime BSI sCMOS camera (Teledyne Photometrics), respectively. Cells were illuminated using CELESTA light engines (Lumencor; NW Greenbrier Parkway Beaverton, OR, USA).

### AMATERAS imaging

Imaging using an AMATERAS-1 system was performed as previously described (Ichimura *et al*, 2021). A CMOS camera (VCC-120CXP1M; CIS, Tokyo, Japan) with a 120-megapixel image sensor (120MXSM; Canon, Tokyo, Japan) was used. A 2× macro lens (LSTL20H-F; Myutron, Tokyo, Japan) was used as the imaging lens. A high-power LED with a center wavelength of 525 nm (SOLIS-525C; Thorlabs, Newton, NJ, USA) was used for fluorescence imaging of O-GECO with an excitation filter (# 86-963; Edmund Optics, Barrington, NJ, USA) and a fluorescence filter (#67-048; Edmund Optics). A stage-top incubator (U-140A; BLAST, Kawasaki, Japan) was used to control the temperature and CO_2_ concentration at the sample stage. The sample stage position and angle were adjusted using three translation stages (TSD-651C25-M6, TSD-651C-M6, and TASB-653-M6; OptoSigma, Tokyo, Japan) and two goniostages (GOHT-65A50BMS-M6 and GOHT-65A75 BMSR-M6; OptoSigma).

### Frequency analysis of iCWPs

The fluorescence intensity of each pixel was measured over time, and the derivatives calculated. Binary images were generated for every time point by identifying pixels with a fluorescence intensity higher than that of the first plane, based on the differential coefficient of each pixel. Next, pixel aggregates having an area > 100 µm^2^ were automatically selected and enclosed with bounding boxes by the ImageJ module Analysis Particles. In consecutive planes, quadrilaterals with a distance ≤ 12 µm between bounding boxes were grouped together as a single population. The areas of the aggregates were measured over time, and a population with a maximum area > 500 µm^2^ (equivalent to 4 cells) was counted as a single wave propagation.

Alternatively, for the images captured using AMATERAS, cell populations with increased fluorescence intensity were detected by Igor Pro v.8.04 (WaveMetrics, Portland, OR, USA), and those with an area > 500 µm^2^ were identified as waves. Next, the images were segmented into 200 × 200 µm square compartments (40,000 µm^2^). The number of waves within each compartment was integrated for 600 s and displayed on a color map.

### Immunofluorescence-based virus infection assay

Immunofluorescence-based viral internalization and infection assays were performed as previously described (Fujioka *et al*, 2013, 2018). In brief, cells were incubated with viruses for 4 h at 37 °C, fixed with 3% PFA for 15 min at 25 °C, and incubated with 1% bovine serum albumin to block nonspecific binding of antibodies. Cells were further incubated overnight at 4 °C with anti-IAV NP antibodies (1:1000 dilution), after which the immune complexes were detected by incubation with Alexa Fluor 488- or Alexa Fluor 647-conjugated secondary antibodies (1:250 dilution) for 1 h at room temperature in the dark. Nuclei were visualized using Hoechst 33342. Images were acquired using a Nikon Ti2 microscope (Nikon, Tokyo, Japan). In some experiments, an ImageJ module Curve Fitting was used to fit the datasets to the Hill equations.

### Dispersity analysis of infected cells

Confluent monolayers of MDCK cells in 35-mm dishes were treated with BPTU or DMSO for 1 h before exposure to IAV. After 30 h of inoculation with IAV, cells were fixed with 3% PFA and subjected to an immunofluorescence-based viral infection assay to identify infected cells. Dispersion was determined by measuring the distance between centers of gravity of the infected cell nuclei as follows: First, the center of gravity of each infected cell nucleus was identified using the ImageJ module Analyze particles. Next, the distance from the center of gravity of each infected cell nucleus to that of the nearest infected cell nucleus was measured (measured values). The frequency of the distances obtained was plotted on a histogram.

Alternatively, cells equal in number to infected cells were assigned random center-of-gravity coordinates and arranged on the *xy*-plane. The distances between their nuclei were measured and plotted in the same manner (calculated values). The median values of distances obtained from actual images and images with randomly allocated cells were compared (Student’s *t*- test).

### Endocytosis assay

To evaluate clathrin-independent endocytosis, MDCK cells plated on collagen-coated glass-bottom dishes (35-mm-diameter; Matsunami Glass Industry Glass, Osaka, Japan) were incubated with Alexa Fluor 546-conjugated dextran (100 μg/mL) for 30 min at 37 °C, followed by thorough washing with PBS to remove noninternalized substances. Visualized vesicles were extracted using the ‘granularity’ function of MetaMorph software (Molecular Devices, San Jose, CA, USA), and the fluorescence intensity of vesicles was quantified.

### Quantification of viral RNA by quantitative PCR (qPCR) analysis

Total RNA was isolated from IAV-infected cells using a QIAamp Viral RNA Mini Kit (Qiagen, Venlo, Netherlands). IAV-infected lung tissues were subjected to total RNA extraction using a PureLink RNA Mini Kit (Thermo Fisher Scientific). Portions of RNA (2.0 μg) was subjected to reverse transcription using SuperScript VILO Reverse Transcriptase (Thermo Fisher Scientific). qPCR analysis was performed using a StepOne real-time PCR system (Thermo Fisher Scientific) with the following primers:

5’-CCMAGGTCGAAACGTAYGTTCTCTCTATC-3’,

5’-TGACAGRATYGGTCTTGTCTTTAGCCAYTCCA-3’, and

probe 5’-FAM-TGACAGRATYGGTCTTGTCTTTAGCCAYTCCA-BHQ1-3’.

### Animal experiments

Six-week-old male BALB/C mice were purchased from Japan SLC Inc. (Shizuoka, Japan). Mice were infected by direct delivery of PR8 (50 PFU) to the nares using a micropipette under isoflurane anesthesia, as previously described.(Fujioka *et al*, 2018) Mice were separately housed for 2 days after virus exposure. BPTU and oseltamivir were dissolved in 5% sucrose solution and adjusted to 16 and 40 mg·kg^-1^·day^-1^, respectively. Mice were intraperitoneally administered these agents or PBS daily for 1 day prior to virus exposure. RNA was extracted from the collected lungs, and viral RNA copy number was determined using qPCR, as described above.

### Quantification and statistical analysis

Quantitative data are presented as mean + standard error (SEM) of at least three independent experiments, except for an animal experiment, and were compared using Student’s *t*-test (parametric test between two conditions) or one-way analysis of variance (ANOVA) followed by Dunnett’s post hoc test (among multiple conditions). No statistical methods were used to predetermine sample size. All experiments were performed unblinded. A *p*-value < 0.05 was considered statistically significant, and all statistical analyses were performed using JMP Pro software v.16.0.0 (SAS Institute Inc., Cary, NC, USA).

## Acknowledgments

We thank Dr. Y. Kawaoka and Dr. K. Aoki for the expression plasmids, viruses, and cells; Dr. T. Mori for advice on a solvent for *in vivo* drug administration; Dr. S. Onami for the transfer of imaging data acquired by AMATERAS; and A. Kikuchi, T. Saeki, and M. Fujioka for technical assistance. This work was supported in part by Grants-in-Aid for Transformative Research Areas (A) (#20H05872) and Scientific Research on Innovative Areas (#15H01248, #26115701, #19H05411, #19H04823, and #21H00413) from the Ministry of Education, Culture, Sports, Science and Technology of Japan, Grant-in-Aid for Scientific Research (B) (#21H02735 and #24K02255) from the Japan Society for the Promotion of Science, Research Program on Emerging and Re-emerging Infectious Diseases (JP20fk0108401), and AMED-CREST (JP20fk0108401) from the Japan Agency for Medical Research and Development, the Photo-Excitonix Project in Hokkaido University, and KONICA MINOLTA Science and Technology Foundation, the Mochida Memorial Foundation for Medical and Pharmaceutical Research.

## Author contributions

Conceptualization: Y.F. and Y.O. Methodology: Y.F., F.K., and Y.O. Formal analysis: F.K., Y.F., N.T., T.K., T.I., and Y.F. Investigation: F.K., T.T., T.K., R.S., Y.H., H.O., and Y.F. Data curation: Y.F. and F.K. Resources: F.K., S.K., and Y.F. Validation: Y.F., F.K., and Y.O. Writing–original draft: Y.F., F.K., and Y.O. Writing–review and editing: F.K., Y.F., T.T., T.K., T.I., S.K., T.K., M.A., T.N., T.F., and Y.O. Visualization: Y.F. and F.K. Supervision: Y.F., M.A., T.N., T.F., and Y.O. Project administration: Y.F. and Y.O. Funding acquisition: Y.F. and Y.O.

## Expanded View Figures Figures EV1-5

**Figure EV1.** iCWPs are ignited by IAV-infected cells, related to Figure 1

**Figure EV2.** Wild-type and mutant IAV M2 are localized to mitochondria, related to Figure 3

**Figure EV3.** The spread rate of iCWPs induced by M2 is comparable to that induced by IAV infection, related to Figure 3

**Figure EV4.** Sustained elevation of Ca^2+^ concentration in M2-expressing cells initiates iCWP, related to Figure 4

**Figure EV5.** M2-induced ICWP facilitates IAV infection in surroundings through enhanced endocytosis, related to Figure 5

**Movies EV1-5**

**Movie EV1.** iCWPs in IAV-infected cells, related to Figure 1A

**Movie EV2.** iCWPs in MOCK-infected cells, related to Figure 1B

**Movie EV3.** iCWPs in IAV-infected cells, as visualized by AMATERAS, related to Figure 2A)

**Movie EV4.** iCWPs in MOCK-infected cells, as visualized by AMATERAS, related to Figure 2B

**Movie EV5.** iCWPs in M2-transfected cells, related to Figure 3A

